# mRNA capping enzyme exports to cytoplasm, localizes to stress granules and maintains cap homeostasis of the target mRNAs

**DOI:** 10.1101/2023.03.28.534620

**Authors:** Anakshi Gayen, Avik Mukherjee, Shubhra Majumder, Chandrama Mukherjee

**Affiliations:** Institute of Health Sciences, Presidency University, Kolkata 700124

**Keywords:** mRNA capping enzyme, Nuclear Export Signal, cytoplasmic capping, stress granules, Xrn1 susceptibility, cap homeostasis

## Abstract

mRNA decapping is believed to trigger RNA degradation until the identification of cytoplasmic capping that has changed the epitome of RNA stability. Unlike nuclear capping machinery that includes RNA polymerase II bound mRNA Capping Enzyme (CE), N-7 RNA methyl transferase and RNMT activating protein RAM, cytoplasmic capping complex consist of cytoplasmic pool of CE (cCE) and N-7 RNA methyl transferase-RAM along with a few cytoplasmic proteins of various functions. Cytoplasmic capping has been shown to recap selective uncapped mRNAs and maintains cap homeostasis by a cyclic process of decapping and recapping. Thus, it acts as post-transcriptional nexus for the target transcripts. Our data show nuclear export of mammalian CE is regulated by Exportin1 (XPO1) pathway via a conserved Nuclear Export Signal sequence. In order to examine biological function of cCE, we show cCE forms granules during stress and majority of these granules co-localize with SGs. In order to identify how cCE regulates cap homeostasis during stress and recovery, we measured the cap status of specific cCE targeted mRNA transcripts along with non-targeted transcripts during non-stress, stress and recovery phase using Xrm1 susceptibility assay. Our data show cCE targeted mRNA transcripts lost their caps in stress condition when cCE is sequestered in granules. After removal of stress, when cCE is released, the cap status has been restored for these transcripts pointing towards the role of cCE in altering cap homeostasis and thus promoting cellular recovery from stress.

## Introduction

Every RNA synthesized by RNA polymerase II contains a 7-methylguanosine ‘Cap’ that is added to the 5’-end of the nascent RNA by the action of mRNA Capping Enzyme (CE) in the nucleus, and is critical to maintain the integrity of mRNAs [1]. This process involves the formation of an unique 5’-5’ triphosphate linkage between a guanosine mono phosphate (GMP) to the first transcribed nucleotide, followed by methylation of the guanosine moiety at N7 position by RNA (N-7) Methyl Transferase (RNMT) thereby synthesizing N7-methylguanosine (m^7^G) [2, 3]. This co-transcriptional process provides mRNA stability, maintains integrity, helps in the transport from nucleus to cytoplasm and in translation initiation [4]. The loss of cap from the transcripts via decapping pathways were thought to be irreversible, leading to decay of all transcripts until the discovery of recapping of the selected transcripts by the cytoplasmic Capping Enzyme (cCE) in mammalian cells [5]. Evidence corroborates the presence of cytoplasmic capping activity in *Drosophila or Trypamosome* as well [6, 7]. While *Trypanosoma* encodes different proteins for nuclear and cytoplasmic capping (CE and cCE respectively), there is only one gene expressing CE homolog in *Drosophila* and mammals, indicating nucleo-cytoplasmic shuttling of that protein operates in these two organisms. Indeed, a recent study demonstrated such shuttling of CE in *Drosophila*, which could be blocked by inhibitor of nuclear export [6]. However, if a similar process is active in mammals is not known yet.

cCE acts in association with adaptor protein Nck1, a complex of RNMT and its activator, RAM, and a yet-to-be-identified 5’-monophosphate RNA kinase [3, 5, 8, 9]. Our previous study discovered cCE to play a role in maintaining decapping and recapping of the target mRNAs, namely ‘cap homeostasis’ [10]. To elucidate specific role of CE restricted to cytoplasm, this study utilized the dominant negative form of CE where the lysine at the active site (K294) was mutated to alanine, a nuclear localization signal (NLS) containing four amino acids (KRKY) at its C-terminal was deleted, and a nuclear export sequence (NES) from human immunodeficiency virus (HIV) was introduced at the N-terminal [5]. [5]. Controlled overexpression of that construct (referred to as K294A) under doxycycline induction in a stable cell line results in the accumulation of uncapped transcripts that were identified to code for proteins associated with nucleotide binding, RNA and protein localization and mitotic cell cycle [10]. Importantly, overexpression of K294A reduced the cell viability upon brief arsenite stress, suggesting that downregulation of cytoplasmic capping activity attenuated cell viability in response to oxidative stress [5].

Stress evolves from both biotic and abiotic conditions culminating in the accumulation of a membrane-less aggregates in the cell termed as stress granules (SGs) [11-13]. These are transiently formed cellular compartments assembled only upon stress and consisting mainly of translation initiation factors and non-translating messenger ribonucleoproteins (mRNPs), which are disassembled after stress removal [14]. The composition and function of SGs generated from diverse stresses may vary. In oxidative stress, this dynamic and reversible nature of SGs promotes cell survival by maintaining homeostasis through RNA and proteins, whereas pro-apoptotic SGs generated by Nitric oxide promote cell death [15].

Here, we discovered that mammalian CE can shuttle between nucleus and cytoplasm, like fly CE. However, this shuttling of mammalian CE is mediated by a novel putative NES in CE, which is conserved among vertebrates, and is dependent on XPO1/Exportin1 pathway for the export of CE to cytoplasm. Moreover, we show here that CE is localized to SGs during brief oxidative stress and cCE-target mRNA transcripts remain in their uncapped states., These mRNA transcripts can be recapped after the removal of stress when cCE is released, suggesting an important role of cCE in cellular stress response by altering cap homeostasis of target mRNAs.

## Materials and Methods

### Constructs

Mammalian expression construct of mouse CE (mCE) pcDNA3-bio-myc-mCE, generated during our previous study[10], was purchased from Addgene (#82475). The pcDNA4/TO-bio-myc-mCE construct was prepared by sub-cloning the bio-myc-mCE sequence into an empty pcDNA4/TO vector using KpnI and ApaI restriction sites. All the mutant constructs of CE were prepared from pcDNA4/TO-bio-myc-mCE by using In-Fusion Site Directed Mutagenesis kit (Takara Bio). mCherry-H2B, GFP-G3BP and Rennila Luciferase constructs were generously provided by Dr. Benubrata Das (IACS, India), Dr. Nancy Kedersha (Harvard Medical School, USA) and Dr. Somshubhra Nath (Presidency University, India).

### Cell culture, transfection, sodium arsenite treatment and heterokaryon fusion assay

Human osteosarcoma cells U2OS (ATCC), Mouse fibroblast NIH-3T3 (Takara), human retinal pigment epithelial cells (hTERT-RPE1 or RPE1, ATCC) were cultured in Dulbecco’s modified eagle’s medium (DMEM) containing 10% fetal bovine serum (FBS) and 1% penicillin-streptomycin mix at 37°C in 5% CO_2_. Jet prime (Polyplus) reagent is used for transfection according to the manufacturer’s instructions. U2OS cells were used for majority of the assays where 1.5×10^5^ cells were seeded in each 35 mm dish. To inhibit nuclear export, cells were treated with 5 and 10 nM of Leptomycin-B (LMB) for 16 hours. For introducing oxidative stress, cells were treated with 0.5 mM sodium arsenite (NaAsO_2_) for 1 h. To recover cells from stress, sodium arsenite-containing medium was replaced with fresh warm media and the cells were incubated for additional 2 h. Interspecies heterokaryon fusion assay was conducted using U2OS and NIH-3T3 cells according to a previously published protocol[16].

### CE knockdown and cell viability assay

For siRNA treatment, 7.5× 10^4^ U2OS cells were seeded in each 35 mm dish and 2×10^3^ U2OS cells were seeded in each well of a 96-well plate. Cells were transfected with 10 nM siRNA against CE (s16640, Invitrogen) or silencer negative control (Ambion) using RNAiMAX (Invitrogen) following manufacturer’s instruction. Cell viability was measured from the cells growing in 96-well plate using XTT assay kit (Cell Signaling Technology) following manufacturer’s instruction using BioTek Synergy microplate reader.

### Nuclear and cytoplasmic fractionation

Following transfection and LMB treatment, U2OS cells were harvested using a cell scraper followed by centrifugation at 1000×g for 5 min. The cell pellet was resuspended in 5 volumes of cytoplasmic lysis buffer (20 mM Tris-HCl, pH 7.5, 100 mM NaCl, 10 mM MgCl_2_, 10 mM KCl, 0.2% NP40 supplemented with 1 mM PMSF and 1X protease inhibitor cocktail) and incubated on ice for 10 min with occasional flicking of the tube. After centrifuging the suspension at 1000xg for 10 min at 4°C, the nuclei were collected as pellet, while the supernatant is transferred to a fresh tube as the cytoplasmic extract. The nuclear pellet was then lysed in a nuclear lysis buffer (25 mM Tris-HCl, pH 7.5, 150 mM NaCl, 1% Triton X-100, 0.5% sodium deoxycholate, 1% SDS supplemented with 1 mM PMSF and 1X protease inhibitor cocktail) that was used half the volume of cytoplasmic lysis buffer, incubating on ice for 45 min with occasional vortexing. Finally, nuclear extract was collected by centrifugation at 10, 000 g for 10 minutes at 4 °C.

### Stress granule isolation

Stress granules were isolated from the cytoplasmic fractions of sodium arsenite treated U2OS cells according to the protocol described in [17], which produced sub-fractions as shown in the schematic diagram (Figure 3). Proteins were isolated from cytoplasmic fractions of depleted SG, enriched. Efficiency of granule isolation was determined by immunoblotting using antibodies against SG-specific proteins. Cytoplasmic extract from unstressed cells was considered as control.

### Western blotting

Cellular protein extracts or immunoprecipitated protein complex was mixed with 1X Laemmli buffer, analyzed on by SGS-PAGE, and western transferred onto Immobilon-FL PVDF membrane (Millipore). Membranes were blocked in 3% Bovine Serum Albumin (SRL) in phosphate-buffered saline (PBS) for 30 min at room temperature, and then incubated with primary antibodies, diluted in the blocking buffer according to the manufacturer’s guidelines, overnight at 4°C. The primary antibodies used here are-rabbit anti-CE (#EPR19384; Abcam), rabbit anti-G3BP1 (#13057-2-AP; Proteintech), rabbit anti-Xrn1 (#PA5-57110; Invitrogen), mouse anti-α Tubulin (#SC-17787; SCBT), goat anti-eIF3 (#SC-16377; SCBT), mouse anti-GAPDH (#2D4A7; Novus), mouse anti-Myc (#32293; SCBT), rabbit anti-KDM1 (#EPR6825; Abcam). After washing in PBS-T, the membranes were incubated in secondary antibodies that were DyLight 800-conjugated donkey anti-mouse (#SA535521; Invitrogen) or Alexa Fluor 680-conjugated donkey anti-Rabbit (#A143; Invitrogen) or anti-Goat (#A11055; Invitrogen) at 1:10000 dilutions in PBS-T. Finally, the immunoblots were scanned and analyzed using Odyssey CLx infrared imaging system (LiCor).

### Immunoprecipitation

Oxidative stress was introduced to U2OS cells grown in 100 mm dish (80-90% confluent) as described above. Cytoplasmic extract prepared from the harvested cells was used for immunoprecipitation (IP) using ProteinG magnetic beads (Biorad). Beads were first equilibrated with IP buffer (25 mM Tris-Cl, pH 7.5, 150 mM NaCl) and then blocked with blocking buffer (1% BSA in IP buffer) for 30 min, followed by incubation with 2 μg anti-CE antibody for 1 hr at room temperature at shaking condition. Next, the antibody bound beads were mixed with the precleared cytoplasmic extract and incubated overnight at 4°C in shaking condition. As control, beads for preclearing the cytoplasmic extract were used. Both control and antibody bound beads were extensively washed with IP wash buffer (0.1% NP40 in IP buffer). The bead bound proteins, and 3% of the input were analyzed by immunoblotting.

### Indirect Immunofluorescence

Cells were grown on coverslips, and were fixed with freshly prepared fixing solution containing 4% paraformaldehyde (EMS) and 0.2% TritonX-100 (Biorad) in 1X PBS for 10 min at room temperature. After washing in 1X PBS containing 0.5mM MgCl_2_ and 0.05% Triton X-100, cells were incubated in a blocking buffer (5% FBS, 0.2 M glycine, 0.1% TritonX-100 in 1X PBS) for 30 min at room temperature. Then the cells were incubated with the primary antibody diluted in the blocking buffer for overnight at 4°C. Primary antibodies used here were-mouse anti-Myc (1:200, SCBT); rabbit anti-eIF3 (1:1000, SCBT); rabbit anti-CE (1:100, Novus); anti-TIA (1:200, SCBT). After washing, the cells were incubated with donkey anti-rabbit, donkey anti-goat or donkey anti-mouse secondary antibodies conjugated with either Alexa Flour 488 or Alexa Fluor 568 (1:1000; Invitrogen), and Hoechst 33358 (Sigma) diluted in the blocking buffer, for 45-60 min at room temperature. For staining the cytoskeleton, phalloidin conjugated with Alexa Fluor 488 or 546 (1:200, Invitrogen) was added to the secondary antibody mix. The coverslips were mounted on glass slides using slowfade mounting media (Invitrogen). Images were acquired using Confocal laser scanning microscope (Leica) fitted with a 63X Plan Apo oil immersion objective (NA 1.4). Images were analyzed, and the fluorescence intensity measurements were performed using associated Las-X software package.

### *In silico* analysis to identify NES

The plausible nuclear export signals of both mouse and human capping enzymes were identified by using online Bioinformatic tool LocNES (available in the URL: http://prodata.swmed.edu/LocNES/LocNES.php). Amino acid sequence of CE that shows the highest score from above analysis is chosen for further verification. As positive control, sequence of human cAMP-dependent protein kinase inhibitor alpha containing XPO1 dependent NES and as a negative control sequence of human histone H1 lacking any XPO1 dependent NES were used for *in silico* analysis.

### *In vitro* Xrn1 susceptibility assay

Fifty nanogram of cytoplasmic poly(A) selected RNAs from three independent experiments was heat denatured and treated with 0.5 units of Xrn1 (New England Biolabs) for 1 h at 37 °C. The RNA was recovered after phenol chloroform extraction followed by ethanol precipitation. RT-qPCR was performed using gene-specific primer pairs against the 5’ ends of the candidate transcripts as done in earlier studies[10, 18, 19]. Here, a synthetic uncapped GFP mRNA and a synthetic capped Renilla luciferase mRNA were used as a positive and negative spike-in control prior to Xrn1 digestion. The C_t_ values for each cCE targeted mRNAs and non-targeted controls were normalized against the C_t_ value of Renilla luciferase obtained from each condition. ΔX or change in Xrn1 susceptibility in cells from unstressed, stressed and stress recovered conditions were obtained by calculating the differences in normalized Ct values of each transcript before and after Xrn1 digestion.

### Cytoplasmic RNA extraction and poly(A) selection

Cytoplasmic RNA was extracted from the cytoplasmic fractions using Trizol reagent (Invitrogen) following the manufacturer’s protocol. RNAs were treated with DNase I (Invitrogen), as directed by the manufacturer and recovered using phenol: chloroform: isoamyl alcohol (SRL) extraction [20]. 2 µg of RNA from each fraction were poly(A) selected using Dynabeads mRNA DIRECT Kit (Invitrogen) as per manufacturer’s instructions.

### RT-qPCR and statistical analysis

Poly(A) selected RNAs were primed with oligo dT and reverse transcribed using a Super Script III First Strand cDNA Synthesis kit (Invitrogen). Primers were designed against selected transcripts from the 5’-ends and spike in controls as done earlier. For qPCR, gene-specific primers as listed in Table S1, were used and qPCR reactions were performed with SsoAdvanced Universal SYBR® Green Supermix (Biorad#172-5271) using CFX connect Real-Time instrument (Biorad). The graphs were generated using Graph-pad Prism V5 software and a student’s two-tailed unpaired t-test was performed to measure the statistical significance.

## Results and Discussion

### Leptomycin B inhibits the nuclear export of CE

Mammalian CE was found only in the nucleus until the identification of cytoplasmic capping [5, 21]. Earlier study has identified cytoplasmic pool of CE in several erythroid and non-erythroid cells expressing epitope tagged constructs of CE [5]. Since overexpression of transgene may cause mis-localization of the overexpressed protein, we decided to confirm the localization using antibody against CE. We have selected mouse fibroblast diploid cell, NIH-3T3, human epithelial diploid cell, RPE1 and primary cells like mouse astrocyte or mouse cardiomyocyte for immunofluorescence study. Our data showed nuclear as well as cytoplasmic localization of CE in these cells (Figure 1A). Actinin antibody is used to distinguish the cardiomyocyte from fibroblast cells which were co-purified during cardiomyocyte isolation. To determine the specificity of anti-CE antibody, we used recombinant protein of His tagged CE to soak the anti CE antibody prior to staining U2OS cells. The signal intensity of CE staining was reduced when soaked antibody was used compared to non-soaked antibody (data not shown) indicating the specificity of the used antibody.

**Figure 1:**
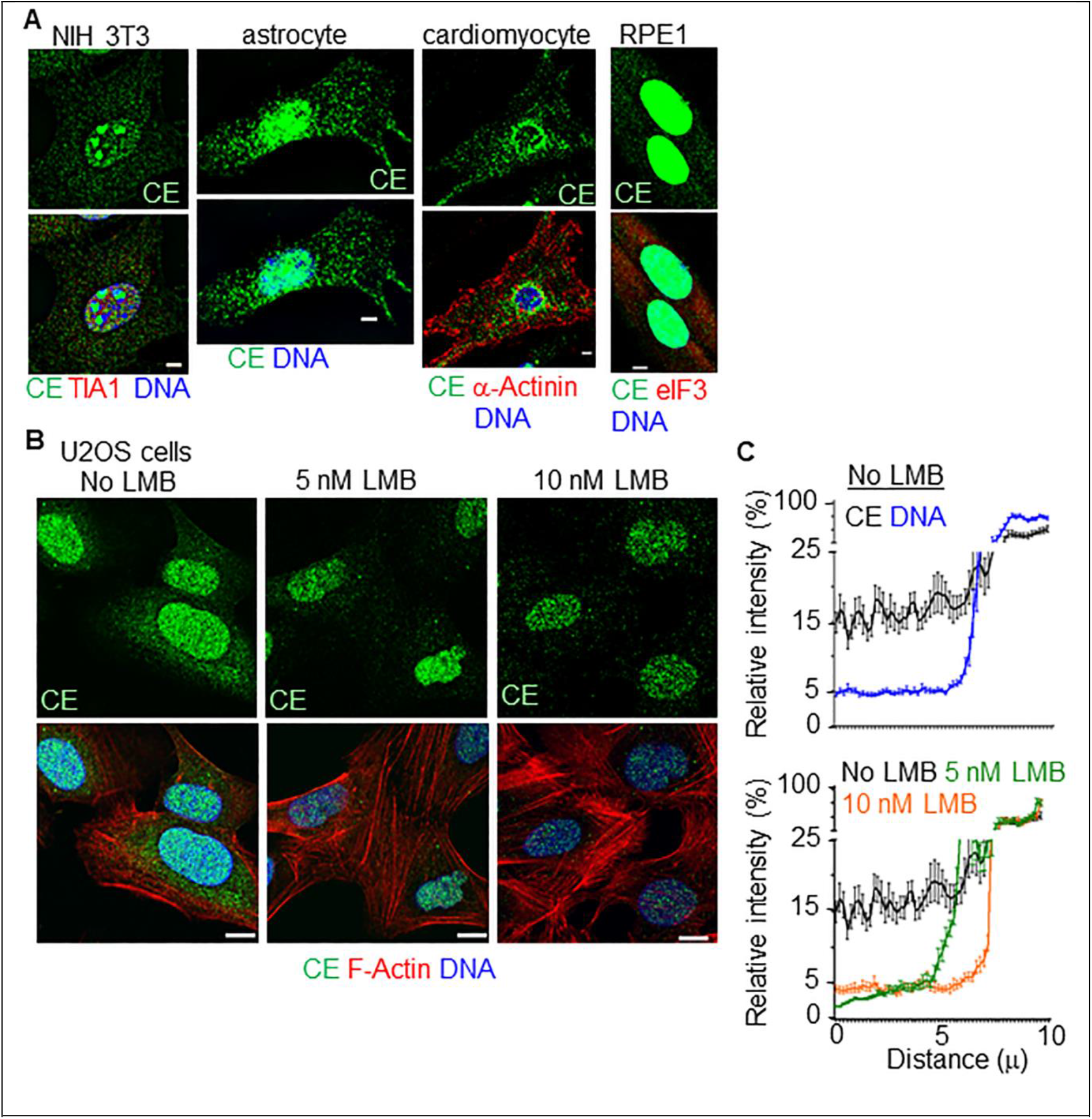
CE is exported to cytoplasm, which is inhibited by Leptomycin B: (A) Representative immunofluorescence images showing the localization of CE (green) at cytoplasm in addition to the prominent staining at nucleus in the indicated mammalian primary cells and diploid cell lines. Indicated antibodies were used for co-staining, bar= 5 µ. (B-C) U2OS cells were treated with different concentration of Leptomycin B (as indicated), and then examined for nuclear and cytoplasmic distribution of CE by quantitative analyses of immunofluorescence images stained for CE. Representative confocal images are shown in B, where phalloidin staining shows the actin cytoskeleton of the cells, which remained undisturbed due to this treatment. Using the line intensity analysis of the image analysis software LasX, fluorescence signal intensities of the indicated fluorophores were measured on a 10 µ line spanning across cytoplasm and nucleus (roughly equal) drawn on the images of 30 randomly chosen cells. (C) The relative intensity profile of CE and DNA staining (upper) and that of CE only (bottom) were plotted in graph where values of each point represent mean ± S.D., n=3 from 30 cells. X-axis represents the distance of that drawn line.

Previous study demonstrated *Drosophila* CE can shuttle between nucleus and cytoplasm [6]. Since Drosophila CE and mammalian CE share sequence similarity, we wondered if mammalian CE can shuttle between nucleus and cytoplasm like drosophila CE. We utilized interspecies heterokaryon fusion with U2OS cells expressing myc tagged mouse CE protein (myc-CE) fused to an equivalent number of NIH 3T3 cells to form interspecies heterokaryons, in presence of cycloheximide to block *de novo* protein synthesis (Supplementary Figure 1A). At the end of the assay, the cells were stained with DAPI to differentiate human and mouse nuclei and examined for the presence of myc-CE protein in the mouse nuclei. Our data showed 40% of mouse cells expressing myc-CE in their nuclei (Supplementary Figure-1A) showing nucleo-cytoplasmic shuttling of CE. Since most of the proteins are exported from the nucleus in XPO1 dependent manner, we aimed to explore if it regulates export of CE.

Therefore, we treated U2OS cells with two different concentrations of Leptomycin B (LMB), a potent inhibitor of XPO1 and stained the cells with antibody against CE. phalloidin to stain actin to mark the individual cell. We found a sharp decrease in cytoplasmic content of CE after 5 nM LMB treatment in comparison to untreated cells (Figure 1B, left and middle panel). 10 nM LMB treatment further reduces the cytoplasmic content of CE (Figure 1B, left and right panel). We have calculated maximum signal intensity of endogenous CE in each cell by drawing a line spanning from cytoplasm to nucleus using Las-X software (see method). DAPI intensity was also calculated in each cell and the percentage of maximum signal intensity was plotted for both DAPI and CE along the line (10 micron) (Figure 1C, upper panel). The plot shows differential distribution of CE, high in nucleus and less in cytoplasm whereas DAPI intensity is only nuclear (Figure 1C, upper panel). LMB treatment yielded a reduction in the maximum signal intensity of cytoplasmic CE from 15% to <5% indicating CRM1/Exportin1 might regulate the export of CE (Figure 1C, lower panel).

### Identification of putative NES in mammalian CE

Next, we examined if LMB treatment showed a similar distribution of CE in cells ectopically expressing Myc-CE. U2OS cells transiently transfected with Myc-CE were treated with 5 and 10 nM LMB. Reduced cytoplasmic staining of Myc-CE was observed after treatment with LMB (Figure 2A, left panel). As a positive control for LMB treatment, we used a modified construct of Myc-CE, used earlier [5] where 12 bp NES sequence from HIV, known to be dependent on CRM1/Exportin 1 pathway, was inserted at the N-terminal and the nuclear localization sequence (NLS) from the C-terminal was deleted. This modified construct known as cCE was used previously to study cytoplasmic capping [5, 8-10, 20, 22]. Cells ectopically expressing Mc-cCE and mcherry-H2B proteins were treated with similar concentrations of LMB. Myc-cCE protein was completely localized in cytoplasm as reported earlier [5] in untreated cells (Figure 2B, right panel). However, LMB treatment inhibited the export of myc-cCE to cytoplasm resulting mostly nuclear localization of the protein (Figure 2B, right panel). mCherry tagged H2B protein, devoid of any NES sequence, acting as the negative control, was independent of LMB treatment and its nuclear localization remained unchanged in LMB treated or untreated cells (Figure 2B, right panel). In order to quantitate the reduction in cytoplasmic staining of Myc-CE or Myc-cCE, we adopted the similar method of calculating maximum signal intensity of fluorescence tags and DAPI from 30 independent cells as stated in the above section. Our analysis showed reduction in the maximum signal intensity of Myc-CE in cytoplasm from 15% to <5% (Figure 2B) like endogenous CE. As expected, Myc-cCE showed maximum reduction in cytoplasmic staining (75% to <10%) and localization mcherry-H2B remained unaffected in LMB treated cells (Figure 2B).

**Figure 2:**
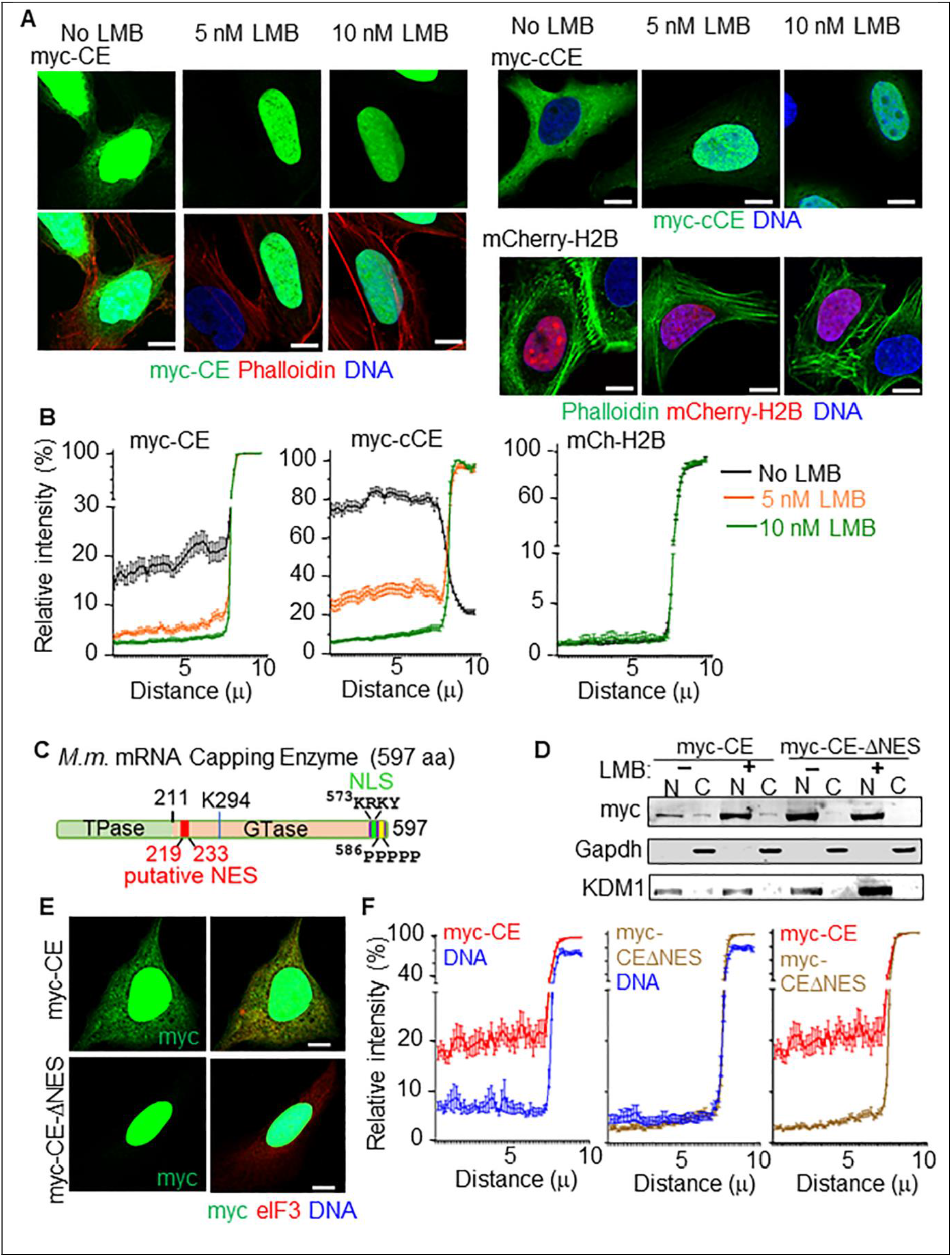
CE uses a putative exportin1-dependent NES motif for export to cytoplasm. (A-B) U2OS cells transfected with plasmid that were ectopically expressing indicated proteins were treated with LMB for 16 h. (A) Representative confocal images show the reduction in cytoplasmic localization of CE due to LMB treatment in a dose dependent manner, bar= 5 µ. Here, myc-cCE that contains a known exportin1-dependent NES serves as a positive control, whereas mCherry-H2B that remains exclusively in nucleus acts as a negative control of the LMB treatment assay. (B) The relative line intensities of the indicated fluorophores were plotted in graph where values of each point represent mean ± S.D., n=3 from 30 cells, similar to the previous figure. (C) Cartoon diagram of mouse mRNA capping enzyme indicating the position of the putative nuclear export sequence (NES) within it. (D) U2OS cells transfected with plasmids expressing myc-CE or myc-CE∆NES were harvested, and their cytoplasmic and nuclear sub-cellular fractions were analyzed by western blotting that show the distribution of these two myc-tagged, ectopically expressed proteins in nuclear and cytoplasmic fractions. GAPDH and KDM1 serve as a loading control for cytoplasmic and nuclear fractions respectively. (E) Representative confocal images show the reduction in nuclear export of myc-CE when the putative NES was removed (myc-CE-ΔNES), bar= 5 µ. (F) The relative line intensities of the myc tag in two cells transfected with myc-CE or myc-CE-ΔNES were plotted in graph where values of each point represent mean ± S.D., n=3 from 30 cells, similar to the previous figure.

**Figure 3.**
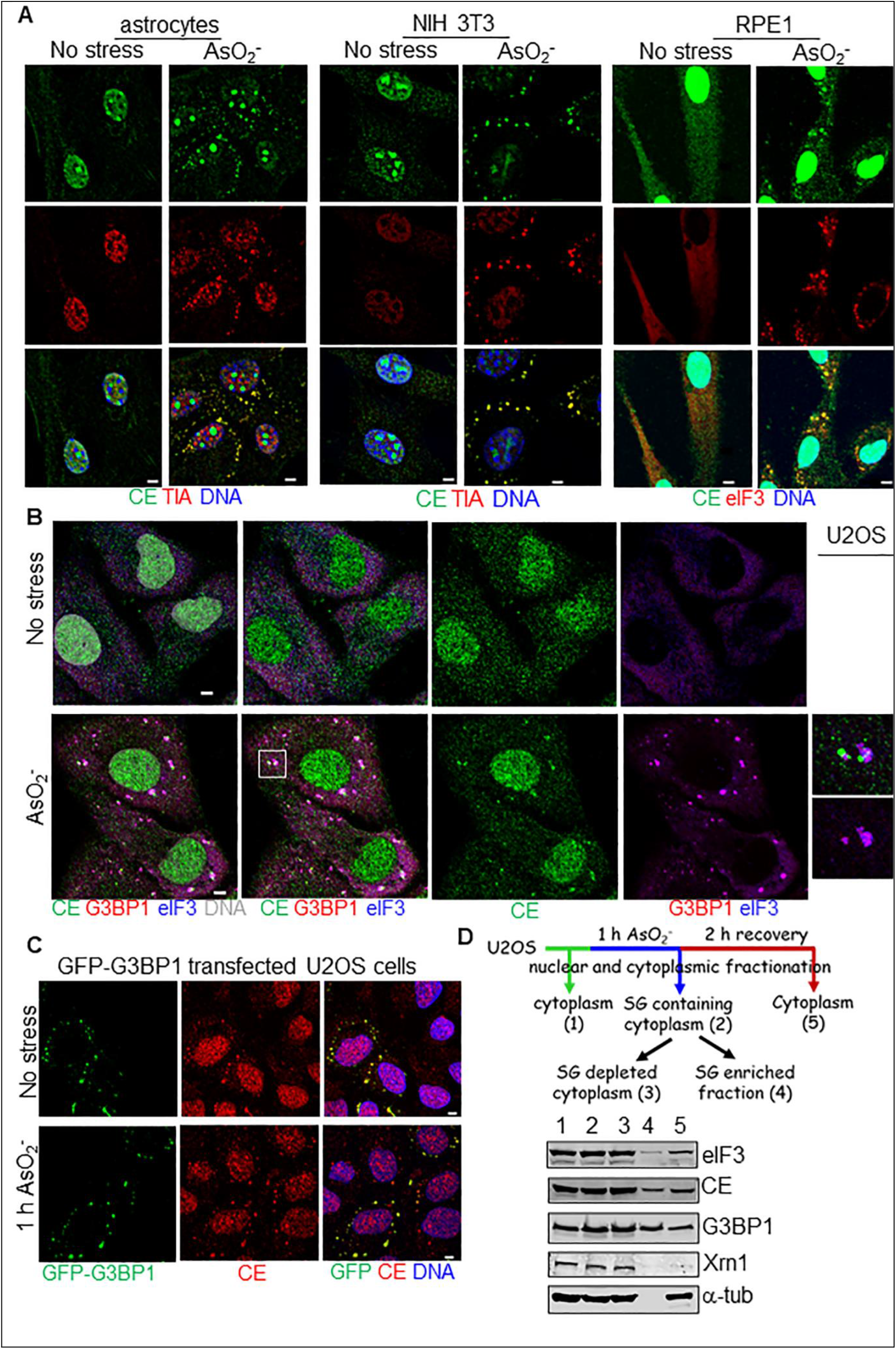
CE is a bona fide candidate of SGs: (A-B) Representative confocal micrographs of indicated mammalian primary cells or cell lines, either untreated or treated with sodium arsenite to introduce oxidative stress, stained for indicated antibodies and DNA. The stress granules containing various SG markers such as eIF3, TIA or G3BP1 were identified in stressed cells. CE that was diffusely localized in the cytoplasm in untreated cells, formed granules upon arsenite stress, majority of which were partially or completely juxtaposed on those eIF3-positive or TIA-positive (A) or eIF3 and G3BP1-positive (B) granules; bar= 5 µ. In B, insets show digitally magnified area surrounded by a square shown on the triple-stained images of the stressed cells. (C) Representative micrographs of transfected U2OS cells overexpressing GFP-G3BP1, which were either untreated or treated with sodium arsenite to introduce oxidative stress. Overexpression of G3BP1 induces SGs even in non-stress condition, and CE co-localizes to these G3BP1 marked granules during both conditions; bar= 5 µ (D) The flow chart shows the procedure of biochemical isolation of cytoplasmic proteins from U2OS cells either untreated or treated with sodium arsenite (AsO_2-_) to introduce oxidative stress, followed by recovery in fresh medium. 1-5 denotes the various cytoplasmic fractions obtained from U2OS cells under different treatment. These fractions were analyzed by immunoblots probed with indicated antibodies. eIF3 and G3BP were used as SG markers and were associated with all fractions, while XRN1, a marker of processing bodies (PBs) and α-tubulin, a cytoplasmic protein were used as negative controls for this assay, which were not detected in SG enriched fraction (4). CE is detected in SG enriched fraction (4) indicating the association of CE with eIF3 and G3BP-positive granules.

Next, to identify CRM1 dependent NES in mammalian CE, LocNES (Locating Nuclear Export Signals or NESs), an online computational tool[23] was employed. We identified a putative NES encompassing amino acids 219-233 in the N-terminus of CE (Figure 2C). To investigate the function of this putative NES, a deletion construct of CE was made lacking amino acids 219-233 with N terminal Myc tag (Myc-CEΔNES) (Figure 2D). Equal amount of nuclear and cytoplasmic fractions from U2OS cells, transiently expressing Myc-CE and Myc-CEΔNES were analysed by western blotting using anti-Myc antibody as shown in figure 2D. Our data showed more nuclear accumulation (lanes 1 and 5, Figure 2D) and less cytoplasmic expression of CE (lanes 2 and 6, figure 2D) in cells expressing Myc-CEΔNES compared to wild type cell expressing Myc-CE suggesting this putative sequence bears the nuclear export function. Efficiency of nuclear and cytoplasmic fractions were examined by using antibodies against nuclear marker KDM1 and cytoplasmic marker Gapdh (Figure 2D). We also treated the wild type and mutant cells with 10 uM LMB and our results showed nuclear accumulation and less cytoplasmic expression of Myc-CE compared to untreated cells (lanes 1 and 3, Figure 2D) whereas nuclear expression of Myc-CEΔNES remained unchanged in LMB treated cells compared to the untreated one (lanes 5 and 7, Figure 2D) indicating participation of NES in CRM1 mediated export. Immunofluorescence data also showed approximate 50% reduction in cytoplasmic localization of CE in cells expressing Myc-CEΔNES compared to wild type cell expressing Myc-CE when stained with antibody against CE (Figure 2E-F). Taken together, our results show mammalian CE exports to the cytoplasm through CRM1/Exportin 1 dependent pathway.

### CE is a bona fide candidate of SGs

Earlier studies pointed the link between cytoplasmic capping and stress [5, 18], we sought to determine the localization of endogenous CE during stress. Cells were introduced to stress by adding sodium arsenite (NaAsO_2_) and localization of endogenous CE was checked in comparison with traditional SG markers TIA, eIF3 in astrocytes and different mammalian cell lines like NIH3T3, RPE1 (Figure 3A). We found that CE formed granules during stress and majority of these granules co-localized with TIA and eIF3 (Figure 3A). SGs are the dynamic membrane-less multi component condensates formed by liquid-liquid phase separation and are known to play a cytoprotective role by maintaining storage of untranslated mRNAs, establishing balance between mRNA degradation and translation [13, 24-26]. We also performed triple staining to confirm localization of CE in U2OS cells with SGs using co-staining with antibodies against eIF3 and another SG marker, G3BP (Figure 3B). Interestingly, we found majority of CE granules either co-localize or juxtaposed with eIF3 and G3BP positive SGs (Figure 3B). Next, to examine if co-localization of CE with SG is dependent on NaAsO_2_ generated stress, we mimicked SG generation in U2OS cells by over expressing GFP tagged G3BP which is an RNA binding protein and a central component of SGs leading SG nucleation [27]. Overexpression of G3BP1 induces SG formation in unstressed condition [27]. Similar to previous studies, we have observed G3BP forms large aggregates during non-stress and stress conditions and CE is co-localized to G3BP formed SGs in both conditions suggesting CE as a bona fide candidate of SGs (Figure 3C).

In order to further characterize, SGs were isolated from U2OS cells after 1 h of stress as shown in the schematic in Fig. 3D, using a published method by Wheeler et al. [17]. Proteins were isolated during different stages of SG isolation, for example, cytoplasmic lysate lacking SG or enriched with SG (Figure 3D). The assembly of SG is also reversible [28]. After removal the stressor, during the recovery phase, SGs disassemble and mRNAs are again dispersed into the cytoplasm and involved in translation [25]. We performed a recovery experiment where NaAsO_2_ was removed after 1 h with fresh medium and cells were kept for 2 h to disassemble the SGs. We found dispersed staining of CE in cytoplasm after recovery similar to non-stress condition as shown in the following section (Figure 4A). Cytoplasmic proteins from non-stress and recovered cells as shown in the schematic (Figure 3D) were isolated and these protein fractions in addition to proteins from SGs were western hybridized with antibodies against CE and SG markers. Our results show recovery of CE in every cytoplasmic fraction with or without SGs (Figure 3D). The SG proteins, eIF3 and G3BP also showed the similar distribution pattern (Figure 3D). We have also used antibodies against P body marker, Xrn1 and α-tubulin for western hybridization to examine the efficiency of SG isolation. Results show presence of both Xrn1 and α-tubulin in all cytoplasmic fractions except SGs (Figure 3D, lane 4). These proteins are not associated with SG as published earlier [29, 30] and absence of these proteins from SG enriched fractions indicate the efficiency of the isolation. Interestingly, we noticed 2 hours recovery from stress is not enough to trigger Xrn1 protein synthesis (Figure 3D). Taken together, our results demonstrated CE forms granules during stress, majority of which either co-localized or juxtaposed to SGs.

**Figure 4:**
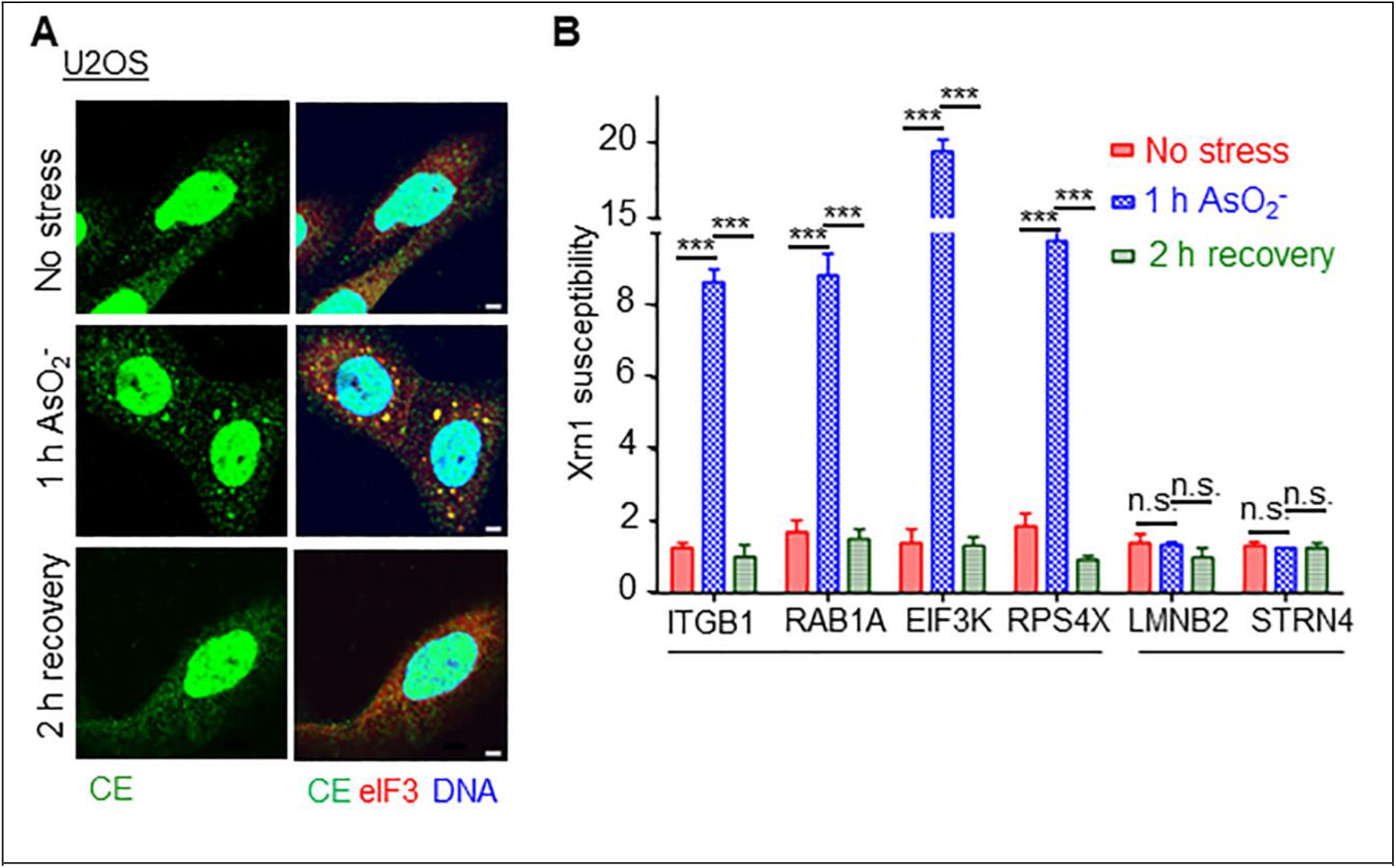
cCE targets are recapped during recovery from stress. (A) Representative confocal micrographs showing localization of CE and eIF3 during non-stress, stress and stress recovered U2OS cells. bar= 5 µ (B) Poly(A)-selected cytoplasmic RNA from three independent sets (n=3, mean ± SD) were extracted from above cells and spiked with capped Renilla luciferase as control RNA. The plot shows Xrn1 susceptibility of each transcript in above conditions where samples were treated with ±Xrn1 followed by RT-qPCR analysis with gene specific primers close to 5’ ends. The susceptibility was measured by the differences in normalized Ct values between two treatment (±Xrn1) groups. For statistical significance two tailed unpaired student’s t-test was performed and asterisks (***) indicate P <0.005.

### cCE recaps target mRNAs during recovery from stress

Inhibiting cytoplasmic capping by overexpressing dominant negative mutant of cCE (K294A) led to poor recovery from arsenite induced stress [5]. Interestingly, another recent study identified TOP mRNAs as recapping targets [18] which has connection with integrated stress responses as observed by single molecule imaging FISH study [31]. Based on these studies, we wondered if cCE has any role in stress response. To address this, we performed *in vitro* Xrn1 digestion that could degrade uncapped RNAs and separate capped and uncapped RNA as done earlier[10, 19]. U2OS cells were fixed before or after addition of NaAsO_2_ and after 2 h recovery from stress. As expected, CE localized to SGs during stress and cytoplasm during recovery or non-stress conditions (Figure 4A). Cytoplasmic poly (A) RNA was isolated from triplicate cultures in these conditions these cells and spiked with capped Rennila luciferase mRNA and uncapped GFP mRNA. Half of the RNA mix were treated with Xrn1 and remaining left untreated followed by RT-qPCR analysis with the gene specific primers close to the 5’ ends. Uncapped GFP mRNA is used to check the efficiency of Xrn1 digestion. The C_t_ values obtained for each transcript in different conditions were normalized against luciferase spike control which remain unchanged with respect to Xrn1 treatment. Xrn1 susceptibility or loss of 5’ ends is represented as the difference in relative C_t_ values before and after Xrn1 digestion. In order to examine role of cCE in recovery, we have examined the cap status of a few recapping mRNA targets (ITGB1, RAB1A, EIF3K, RPS4X) which were validated earlier [10, 18] and control mRNAs (LMNB2, STRN4) which were independent of cytoplasmic capping [10, 18]. We observed the recapping transcripts were more prone to Xrn1 digestion during stress compared to non-stress condition. However, during recovery, these transcripts were less susceptible to Xrn1 (Figure 4B). The control mRNAs did not show any significant change in Xrn1 susceptibility during non-stress, stress or recovery (Figure 4B). Taken together, data presented in Figure 4 showed cCE targets lost their caps during stress when cCE is sequestered mostly in SGs. After withdrawal of stress, during recovery, cCE is released in the cytoplasm and the target mRNAs are less sensitive to Xrn1 digestion. Our data indicate target mRNAs could be recapped by cCE during recovery from stress and thus cCE act in maintaining cap status of the target mRNAs. Overall, our results provide the first evidence showing role of cCE mediated cap homeostasis of target mRNAs in cellular stress response.

## Acknowledgements

The authors sincerely thank Drs Daniel R Schoenberg and Harold Fisk, Ohio State University for providing the lab space to initiate the work on granules, and the access to Fluorescence microscope, respectively. The authors thank Drs Peter Mohler and Stephen Kolb, Ohio State University, USA for providing the primary cells, Dr. Nancy Kedersha, Harvard Medical School, USA, Dr. Benubrata Das, IACS, Kolkata, and Dr. Somsubhra Nath, Presidency University for sharing plasmid constructs. The authors acknowledge financial supports as research grants from Science and Engineering Research Board (SERB), Government of India to CM and SM, Department of Science & Technology and Biotechnology, Government of West Bengal to CM, and Ramalingaswami re-entry fellowship, Department of Biotechnology, Government of India to CM and SM. The authors also acknowledge American Heart Association Fellowship to CM, which helped to procure the reagents to initiate this study. The authors thank Dr. Abhik Saha for providing access to Li-Cor IR imaging system, and Institute of Health Sciences, Presidency University for departmental instrument facility.

## Figures

**Supplementary Figure 1:**
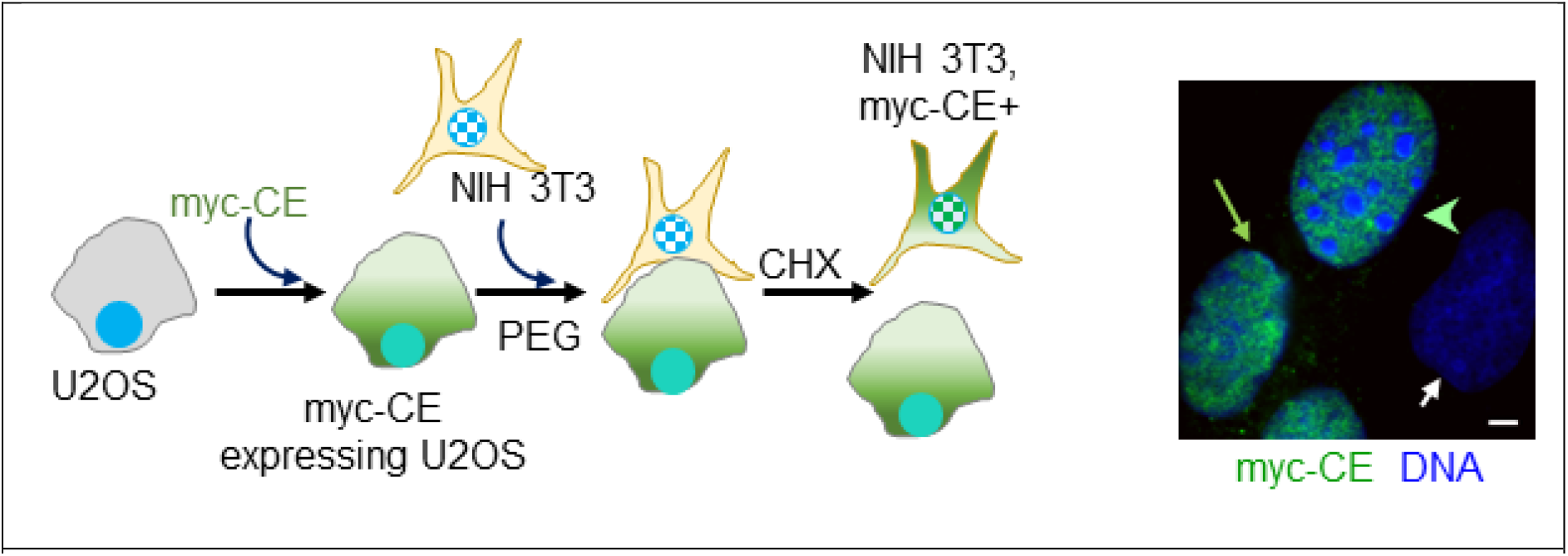
U2OS cells transfected with myc-CE were fused with NIH-3T3 cells and fused with Polyethelene glycol (PEG) andexposed to cycloheximide. The localization of myc-CE was monitored by indirect Immunofluorescence images. CHX-cyclohexamide, Bar represents 5 microns. Arrow indicates U2OS nuclei transfected with myc-CE, arrowhead indicates NIH-3T3 nuclei expressing mcy-CE and small white arrow indicates untransfected nuclei of U2OS cell.

## Notes

### Competing Interest Statement

The authors have declared no competing interest.

## Reference

1. Shatkin, A. J. (1976) Capping of eucaryotic mRNAs, Cell. 9, 645–53.

2. Ensinger, M. J. & Moss, B. (1976) Modification of the 5’ terminus of mRNA by an RNA (guanine-7-)-methyltransferase from HeLa cells, J Biol Chem. 251, 5283–91.

3. Gonatopoulos-Pournatzis, T., Dunn, S., Bounds, R. & Cowling, V. H. (2011) RAM/Fam103a1 is required for mRNA cap methylation, Mol Cell. 44, 585–96.

4. Schoenberg, D. R. & Maquat, L. E. (2012) Regulation of cytoplasmic mRNA decay, Nat Rev Genet. 13, 246–59.

5. Otsuka, Y., Kedersha, N. L. & Schoenberg, D. R. (2009) Identification of a cytoplasmic complex that adds a cap onto 5’-monophosphate RNA, Mol Cell Biol. 29, 2155–67.

6. Chen, P., Zhou, Z., Yao, X., Pang, S., Liu, M., Jiang, W., Jiang, J. & Zhang, Q. (2017) Capping Enzyme mRNA-cap/RNGTT Regulates Hedgehog Pathway Activity by Antagonizing Protein Kinase A, Sci Rep. 7, 2891.

7. Ignatochkina, A. V., Takagi, Y., Liu, Y., Nagata, K. & Ho, C. K. (2015) The messenger RNA decapping and recapping pathway in Trypanosoma, Proc Natl Acad Sci U S A. 112, 6967–72.

8. Mukherjee, C., Bakthavachalu, B. & Schoenberg, D. R. (2014) The cytoplasmic capping complex assembles on adapter protein nck1 bound to the proline-rich C-terminus of Mammalian capping enzyme, PLoS Biol. 12, e1001933.

9. Trotman, J. B., Giltmier, A. J., Mukherjee, C. & Schoenberg, D. R. (2017) RNA guanine-7 methyltransferase catalyzes the methylation of cytoplasmically recapped RNAs, Nucleic Acids Res. 45, 10726–10739.

10. Mukherjee, C., Patil, D. P., Kennedy, B. A., Bakthavachalu, B., Bundschuh, R. & Schoenberg, D. R. (2012) Identification of cytoplasmic capping targets reveals a role for cap homeostasis in translation and mRNA stability, Cell Rep. 2, 674–84.

11. Anderson, P., Kedersha, N. & Ivanov, P. (2015) Stress granules, P-bodies and cancer, Biochim Biophys Acta. 1849, 861–70.

12. Aulas, A., Fay, M. M., Szaflarski, W., Kedersha, N., Anderson, P. & Ivanov, P. (2017) Methods to Classify Cytoplasmic Foci as Mammalian Stress Granules, J Vis Exp.

13. Ivanov, P., Kedersha, N. & Anderson, P. (2019) Stress Granules and Processing Bodies in Translational Control, Cold Spring Harb Perspect Biol. 11.

14. Hofmann, S., Kedersha, N., Anderson, P. & Ivanov, P. (2021) Molecular mechanisms of stress granule assembly and disassembly, Biochim Biophys Acta Mol Cell Res. 1868, 118876.

15. Aulas, A., Lyons, S. M., Fay, M. M., Anderson, P. & Ivanov, P. (2018) Nitric oxide triggers the assembly of “type II” stress granules linked to decreased cell viability, Cell Death Dis. 9, 1129.

16. McNicoll, F. & Muller-McNicoll, M. (2018) A Quantitative Heterokaryon Assay to Measure the Nucleocytoplasmic Shuttling of Proteins, Bio Protoc. 8, e2472.

17. Wheeler, J. R., Jain, S., Khong, A. & Parker, R. (2017) Isolation of yeast and mammalian stress granule cores, Methods. 126, 12–17.

18. Del Valle Morales, D., Trotman, J. B., Bundschuh, R. & Schoenberg, D. R. (2020) Inhibition of cytoplasmic cap methylation identifies 5’ TOP mRNAs as recapping targets and reveals recapping sites downstream of native 5’ ends, Nucleic Acids Res. 48, 3806–3815.

19. Mukherjee, A., Islam, S., Kieser, R. E., Kiss, D. L. & Mukherjee, C. (2023) Long non-coding RNAs are substrates for cytoplasmic capping enzyme, FEBS Lett.

20. Kiss, D. L., Oman, K. M., Dougherty, J. A., Mukherjee, C., Bundschuh, R. & Schoenberg, D. R. (2016) Cap homeostasis is independent of poly(A) tail length, Nucleic Acids Res. 44, 304–14.

21. Wen, Y., Yue, Z. & Shatkin, A. J. (1998) Mammalian capping enzyme binds RNA and uses protein tyrosine phosphatase mechanism, Proc Natl Acad Sci U S A. 95, 12226–31.

22. Kiss, D. L., Oman, K., Bundschuh, R. & Schoenberg, D. R. (2015) Uncapped 5’ ends of mRNAs targeted by cytoplasmic capping map to the vicinity of downstream CAGE tags, FEBS Lett. 589, 279–84.

23. Xu, D., Marquis, K., Pei, J., Fu, S. C., Cağatay, T., Grishin, N. V. & Chook, Y. M. (2015) LocNES: a computational tool for locating classical NESs in CRM1 cargo proteins, Bioinformatics. 31, 1357–65.

24. Ivanov, P., Kedersha, N. & Anderson, P. (2011) Stress puts TIA on TOP, Genes Dev. 25, 2119–24.

25. Riggs, C. L., Kedersha, N., Ivanov, P. & Anderson, P. (2020) Mammalian stress granules and P bodies at a glance, J Cell Sci. 133.

26. Mukherjee, N. & Mukherjee, C. (2021) Germ cell ribonucleoprotein granules in different clades of life: From insects to mammals, Wiley Interdiscip Rev RNA. 12, e1642.

27. Tourrière, H., Chebli, K., Zekri, L., Courselaud, B., Blanchard, J. M., Bertrand, E. & Tazi, J. (2003) The RasGAP-associated endoribonuclease G3BP assembles stress granules, J Cell Biol. 160, 823–31.

28. Anderson, P. & Kedersha, N. (2006) RNA granules, J Cell Biol. 172, 803–8.

29. Kedersha, N. & Anderson, P. (2007) Mammalian stress granules and processing bodies, Methods Enzymol. 431, 61–81.

30. Chernov, K. G., Barbet, A., Hamon, L., Ovchinnikov, L. P., Curmi, P. A. & Pastré, D. (2009) Role of microtubules in stress granule assembly: microtubule dynamical instability favors the formation of micrometric stress granules in cells, J Biol Chem. 284, 36569–36580.

31. Wilbertz, J. H., Voigt, F., Horvathova, I., Roth, G., Zhan, Y. & Chao, J. A. (2019) Single-Molecule Imaging of mRNA Localization and Regulation during the Integrated Stress Response, Mol Cell. 73, 946–958.e7.

